# Polyploid plants have faster rates of multivariate climatic niche evolution than their diploid relatives

**DOI:** 10.1101/406314

**Authors:** Anthony E. Baniaga, Hannah E. Marx, Nils Arrigo, Michael S. Barker

## Abstract

Whole genome duplication is an important evolutionary process in plants. In contrast to other speciation mechanisms, polyploid species begin with substantial postzygotic reproductive isolation from progenitors while being sympatric with one or both. These nascent polyploid species often go extinct due to ecological and evolutionary genetic obstacles. Interestingly, polyploid species appear to quickly occupy different geographic distributions and ecological niches than their diploid progenitors. Using biogeographic data from polyploid and diploid species representing 49 genera of vascular plants, we tested whether climatic niches of polyploid species evolve faster than their diploid relatives. We found polyploid species often have less climatic overlap than expected with diploid progenitors. Consistent with this pattern, we estimated that the climatic niches of polyploid plants consistently evolved faster than the niches of diploid relatives. Our results indicate ecological niche differentiation is important for polyploid establishment, and suggest ecological differentiation is important for speciation processes more widely.

**Statement of Authorship:** AB and MS conceived of project, AB and NA generated the dataset, AB and HM performed analyses, AB and MS cowrote manuscript.

**Data Accessibility Statement:** Upon acceptance all necessary R scripts, data, and files supporting the results will be archived on FigShare with the data DOI included at the end of the article.

## Introduction

The frequency of speciation by polyploidy or whole genome duplication (WGD) has long led to debates concerning their ultimate ecological and evolutionary importance. Among vascular plants, polyploidy is associated with an estimated 15–30% of speciation events (Wood et al. 2009). However, most nascent polyploid lineages have lower estimated net diversification rates than their diploid relatives (Mayrose et al. 2011), possibly due to multiple ecological and evolutionary obstacles (Otto & Whitton 2000; Comai 2005; Arrigo & Barker 2012; Barker et al. 2016a). Genomic analyses reveal that many polyploid species persisted and occur at key locations in the vascular plant phylogeny, such as the ancestry of seed plants (Jiao et al. 2011; Li et al. 2015), angiosperms (Amborella Genome Project), core eudicots (Tuskan et al. 2006; Jaillon et al. 2007; Jiao et al. 2012; Vekemans et al. 2012), as well as taxonomically rich clades like the Asteraceae (Barker et al. 2008; Barker et al. 2016b; Huang et al. 2016; Badouin et al. 2017) and Poaceae (Paterson et al. 2004; Paterson et al. 2009; Estep et al. 2014; McKain et al. 2016). Reconciling these micro– and macroevolutionary patterns of polyploid speciation requires a better understanding of the ecology of polyploid speciation (Barker et al. 2016a; Soltis et al. 2016) that builds on ecological investigations of polyploids in extant taxa (Stebbins 1985; Buggs & Pannell 2007; Munzbergova 2007; Ramsey 2011).

Competition significantly influences the ecological niches of species (Darwin 1859; Gause 1934; Hutchinson 1953; Andrewartha & Birch 1954; Connell 1961; MacArthur 1972). Considering their abrupt origins, interspecific competition likely plays an important role in polyploid speciation because the niches of closely species tend to be more similar to each other than to those of more distantly related ones (Harvey & Pagel 1991; Wiens 2004; Pyron et al. 2015). Newly formed polyploid species are initially imbued with substantial postzygotic isolation from their progenitors, while also sympatric with either one or both parental species (Ramsey & Schemske 1998, 2002). Sympatry with their diploid progenitors is a large obstacle for polyploid establishment because they experience direct competition with either one or both parental species while starting from initially low population sizes (Levin 1975). Mathematical models indicate that polyploid establishment is promoted by high selfing rates, high rates of polyploid formation, local propagule dispersal, and ecological niche differentiation (Levin 1975; Fowler & Levin 1984; Felber 1991; Rodriguez 1996; Husband 2000; Baack 2005; Rausch & Morgan 2005; Fowler & Levin 2016). The importance of ecological niche differentiation is also supported by species coexistence theory (Tilman 1982, 1985; Chesson 2000, 2004) which suggests that coexistence of related species, such as polyploid and progenitor species, is possible if they have different resource needs or utilization strategies.

Some polyploid species are long known to have different geographical distributions, novel ecological niches, and wider niche breadths than their progenitors (Hagerup 1932; Tischler 1937; Wulff 1937; Love & Love 1943; Clausen et al. 1945; Stebbins 1950; Stebbins 1971; Levin 1975). However, previous analyses have found mixed support for the importance of ecological niche differentiation to polyploid species (Stebbins 1971; Felber-Girard et al. 1996; Petit & Thompson 1999; Martin & Husband 2009; Glennon et al. 2014; Marchant et al. 2016). This is surprising given that polyploids may adapt faster than diploids (Orr & Otto 1994; Otto & Whitton 2000; Otto 2007; Selmecki et al. 2015).

To better understand the role of ecological niche differentiation during polyploid speciation, we evaluated the rates of climatic niche evolution of polyploids and their diploid relatives. We explored two aspects of climatic niche evolution in polyploid species. First, we analyzed the amount of climatic niche overlap between allopolyploid species and their diploid progenitors in 25 genera of plants. We then examined whether polyploid species evolved multivariate climatic niches and niche breadths at faster rates than their diploid relatives in 33 genera of plants. We hypothesized that if climatic niche divergence is not important for polyploid establishment, then the rates of climatic niche evolution of polyploid species would not be significantly different than those of diploid species. Conversely, if climatic niche divergence is important for polyploid establishment, we expected polyploid species to have faster rates of climatic niche evolution than related diploids. Our results provide insight into the importance of ecological divergence for polyploid species establishment, and highlight the role of ecological divergence in speciation processes generally.

## Materials & Methods

### Climatic niche overlap between allopolyploid species and their diploid progenitors

Data on allopolyploids and their known parents were collected from the literature. We used the Taxonomic Name Resolution Service v4.0 (Boyle et al. 2013) to verify taxonomy and filter non-valid species. This filtering left 53 trios comprised of two diploid parents and an allopolyploid yielding 132 unique taxa from 16 families and 25 genera. All unique georeferenced locations were sourced and downloaded from the union of the Global Biodiversity Information Facility (GBIF.org), the Consortium of California Herbaria (http://ucjeps.berkeley.edu/consortium/), and the Southwest Environmental Information Network (SEINet). Unique records were retained for each taxon in decimal degrees and kept at a minimum resolution of two decimal points.

### Estimating and Comparing Niche Overlap

We used Ecospat to quantify the multivariate climatic niche overlap among the taxa of a trio (Broennimann et al. 2012). Background areas were defined by adding one decimal degree to a taxon’s maximum and minimum known geographic coordinates to include geographic areas that may have experienced dispersal (Barve et al. 2011). All nineteen current bioclimatic variables 1960–1990 (Hijmans et al. 2005) at 2.5 minute resolution were used to estimate Schoener’s D (Schoener 1970; Warren et al. 2008), a pairwise metric of climatic niche overlap ranging from no overlap (D=0) to complete overlap (D=1). Pairwise analyses of species in each trio were run with 100 iterations and at a 100 pixel resolution.

After pairwise calculations of Schoener’s D for each trio (allopolyploids + known parents), trios were grouped into three different classes of climatic niche overlap. These classes were defined by the amount of climatic niche overlap that an allopolyploid species shares with its parents relative to the amount of climatic niche overlap shared between parents. These include relationship 1 (represented by “P DD”): the allopolyploid species has less overlap with both parents than parents to parents; relationship 2 (“PD D”): the allopolyploid species has less overlap with one parent than parents to parents; and relationship 3 (“DPD”): the allopolyploid species has more overlap with both parents than parents to parents. A two-tailed binomial exact test was used to assess if the observed distribution of these relationships was different than our null expectation of equal frequencies (*p*=0.333).

To examine the association of allopolyploid species age with the amount of climatic niche overlap with its diploid parents, we collected sequences for each taxon and compared the amount of pairwise sequence divergence to pairwise climatic niche overlap. We downloaded available nucleotide sequences of the *trnL*-*trnF* intergenic spacer from PhyLoTA (Sanderson et al. 2008), performed pairwise alignments with MUSCLE v3.8 (Edgar 2004), and inferred pairwise sequence divergence using a GTR + Y nucleotide substitution model in FastTree v2.1.1 (Price et al. 2010).

### Rates of climatic niche evolution in plant polyploid and diploid species

We developed a database of vascular plant polyploid and diploid species including their chromosome numbers (Barker et al. 2016c). We filtered the database for genera with > 20 taxa with known chromosome counts. This filtering left a total of 33 genera from 20 different families comprising 1706 taxa of which 537 were listed as polyploid species. Eight of the allopolyploid comparisons from the section above were also represented here.

For each taxon, all unique georeferenced locations were downloaded from GBIF at a minimum resolution of two decimal points. Six current bioclimatic variables 1960–1990 (Hijmans et al. 2005) important to defining a species climatic niche (BIO1 Annual Mean Temperature, BIO5 Max Temperature of Warmest Month, BIO6 Min Temperature of Coldest Month, BIO12 Annual Precipitation, BIO16 Precipitation of Wettest Quarter, BIO17 Precipitation of Driest Quarter) at 30 arcsecond resolution were extracted for each locality using the R package ‘dismo’ (Hijmans et al. 2016). These bioclimatic variables were chosen to standardize possible variables across species, and because they highlight climatic averages as well as extremes important to defining a species climatic niche (Kozak & Wiens 2006). The arithmetic mean and breadth (i.e., variance) of the six bioclimatic variables were calculated for each taxon, standardized by subtracting the mean followed by division by the variance, and then log transformed. The principal component (PC) loading scores for axes one (PC1) and two (PC2) of the six bioclim variables for each species were calculated using R prcomp and were analysed as continuous traits in OUwie (Beaulieu et al. 2012).

An appropriate outgroup for each genus was found by using the closest BLAST hit to the ingroup. Genus level phylogenies were inferred with MrBayes v3.1.2 (Huelsenbeck & Ronquist 2001; Ronquist & Huelsenbeck 2003) with either a plastid or nuclear molecular marker downloaded from GenBank (see Table S1 in Supporting Information for all comparisons). This molecular dataset is separate from that used to investigate the relationship of allopolyploid age and climatic niche overlap above. To infer changes in ploidal level, we analyzed the highest posterior probability tree for each genus with chromEvol v1 (Mayrose et al. 2010) using available chromosome counts of taxa for each genus (see Table S1).

We accounted for topological uncertainty in our analyses by using the top 50 trees from the posterior distribution of the MrBayes output. Each of the 50 trees were ultrametricized with the chronos function in ape (Paradis et al. 2004), and ploidal shifts were mapped using SIMMAP (Revell 2012). For each of the four multivariate climatic niche traits (PC1–2 niche mean and PC1–2 niche breadth), we used OUwie (Beaulieu et al. 2012) to assess if polyploid and diploid species had different (BMS) or the same (BM1) rates of multivariate climatic niche evolution. We selected the best fitting model using two approaches: 1) a likelihood ratio test (LRT) with an alpha of 0.05 and 2) the corrected Akaike Information Criterion (AICc) where the more sophisticated model (i.e., BMS) was preferred if **Δ**AICc > 4.

For each model test, a relative rate of multivariate climatic niche was calculated by dividing the rate for polyploid species by the rate for diploids when the BMS model was better supported; in instances when the BM1 model was preferred, polyploid and diploid species had the same inferred rate of change and we set the relative rate = 1. The rate of evolution for each trait was summarized by calculating the geometric mean of all 50 model tests and designating each multivariate climatic niche trait as higher, lower, or no difference in rate between polyploids and diploids. We used a two-tailed binomial exact test to assess whether the number of genera observed having a higher, lower, or no difference in rate between polyploid and diploid species was significantly different from chance (*p*=0.333).

To assess the power of our analyses, we compared the observed patterns to null simulations of species distributions. Simulations were conducted for each genus and each trait by randomly shuffling which taxa were assigned as a polyploid, weighted by the number of taxa that were polyploid in that genus, and performing the OUwie model test 100 times per tree for a total of 5000 simulations per trait. Model selection was performed as above with selection by a LRT and AICc. Each model test was binned into a rate class where polyploids could have higher, lower, or no difference in the rates of multivariate climatic niche evolution compared to diploids. We used these frequencies to calculate expected values for a chi-square test (*p*<0.05, df=2) of whether the observed values deviated from the null expectation of equal frequencies of the three rate classes.

## Results

### Climatic niche overlap between allopolyploid species and their diploid progenitors

Our analyses supported the hypothesis that polyploid species have different climatic niches than their diploid progenitors. Of the 53 comparisons involving one allopolyploid species and its two diploid progenitors, 28 (51%) had a “P DD” pattern where the allopolyploid species had less climatic niche overlap with either parent than the parents had with each other (Figure 1). We observed a “PD D” pattern in 12 (23%) cases where the allopolyploid species had less climatic niche overlap with one parent than parents to parents. Finally, 14 (26%) cases had a “DPD” pattern where the allopolyploid species had more climatic niche overlap with both parents than the parents had with each other. Assuming an equal probability of each of these three patterns, we found a statistically significant excess of “P DD” (*p*< 0.01) relative to the other two patterns (Table 1, see Table S2 for all comparisons).

**Table 1.** Summary of allopolyploid trios in each climatic niche overlap scenario. Significance values were calculated with a two-tailed binomial exact test assuming an equal probability of each scenario (*p*=0.333).

|  | Class 1 ‘P DD’ | Class 2 ‘PD D’ | Class 3 ‘DPD’ |
| --- | --- | --- | --- |
| Number of Trios | 27* | 12 | 14 |

*\*(p<0.05)*

**Figure 1.**
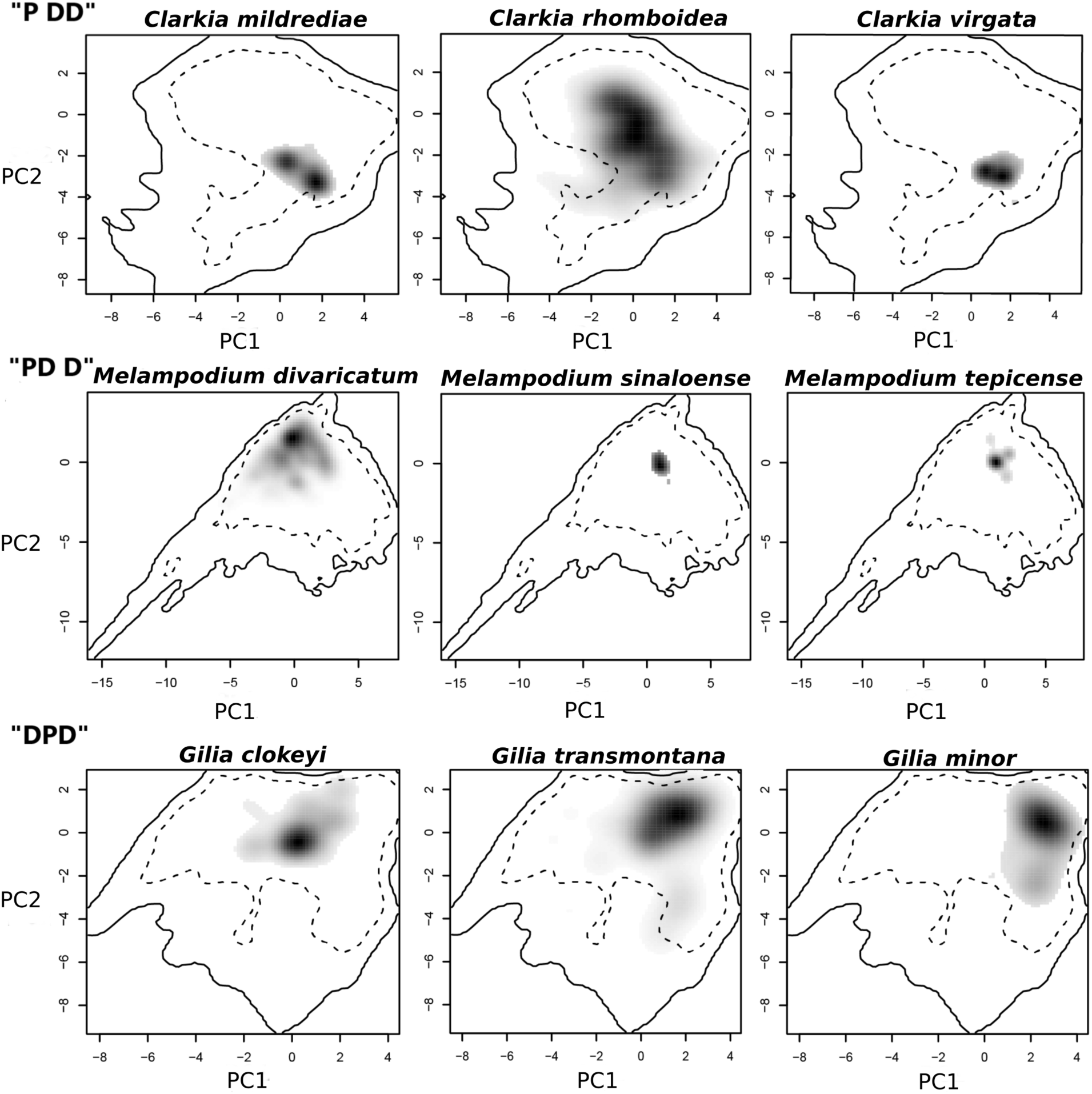
Climatic niche overlap in allopolyploid species and their diploid progenitors. Examples of three different classes of climatic niche overlap, defined by the amount of climatic niche overlap an allopolyploid species shares with its parents relative to that found between parents. Climatic niche overlap diagrams produced by R ecospat (Broennimann et al. 2012). Parental diploid species are represented on the left and right, allopolyploid species in the middle. Examples include: Example 1 (‘P DD’) the allopolyploid species has less overlap with both parents than parents to parents; Example 2 (‘PD D’) the allopolyploid species has more overlap with both parents than parents to parents; Example 3 (“DPD”) the allopolyploid species has more overlap with both parents than parents to parents.

Of the 53 trios of species available to examine the relationship between climatic niche overlap and sequence divergence, 36 species pairs from 20 trios had a suitable *trnL*-*trnF* intergenic spacer available (see Table S3). Pairwise sequence divergence ranged from a low of 0.0 between *Dryopteris corleyi* – *Dryopteris aemula*, and *Lathyrus venosus* – *Lathyrus palustris*, to a high of 0.072 between *Asplenium montanum* – *Asplenium platyneuron* with an average of 0.022. When trios from all three scenarios of climatic niche overlap were examined there was no clear relationship between the amount of climatic niche overlap and sequence divergence (y=2.063x + 0.247; *p*=0.5097, adj-R^2^=-0.161; Figure 2).

**Figure 2.**
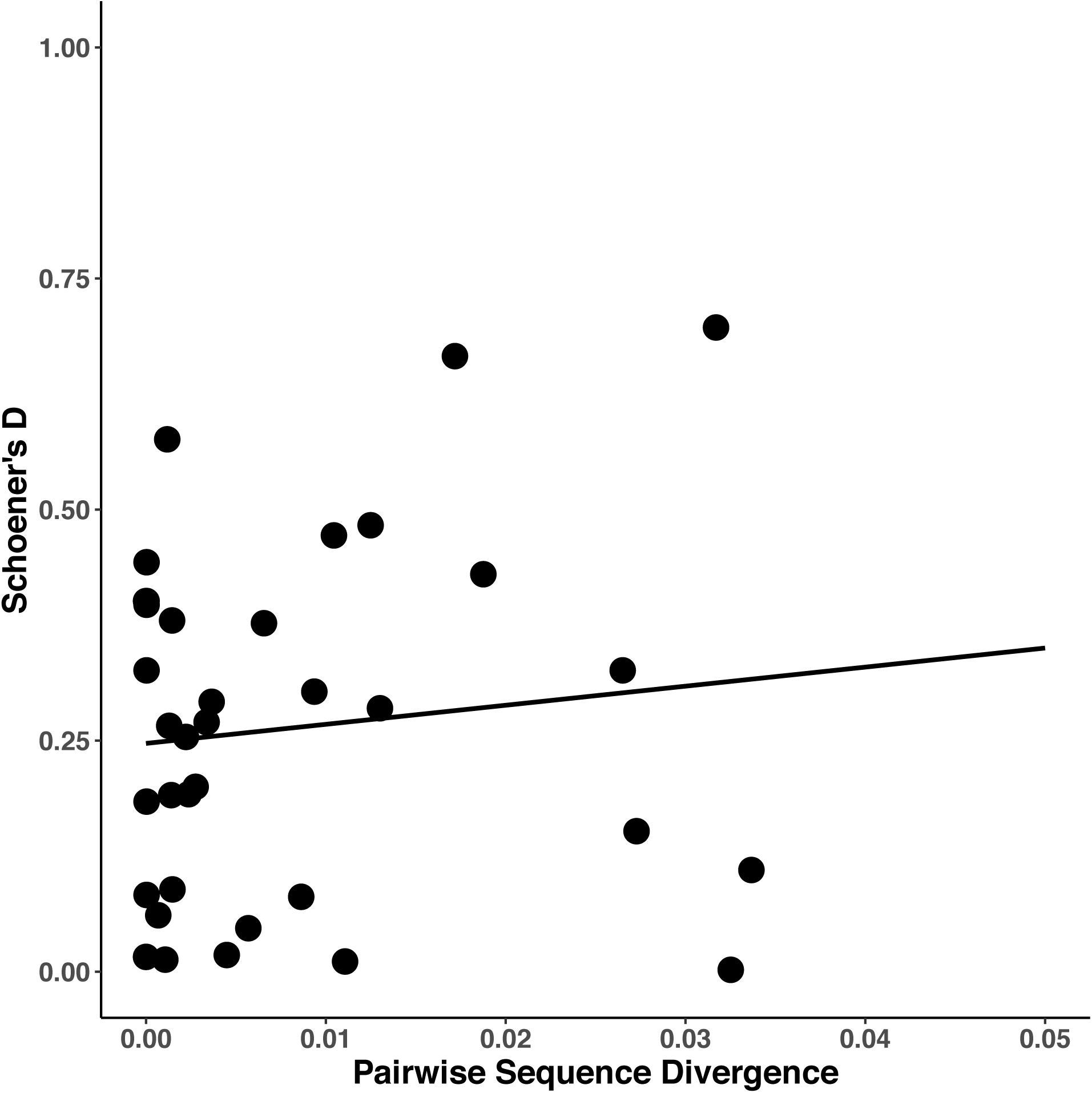
Climatic niche overlap and pairwise sequence divergence. The amount of climatic niche overlap (Schoener’s D) by pairwise sequence divergence estimated from the plastid *trnL-trnF* intergenic spacer. Black circles represent comparisons between an allopolyploid species and one of its diploid parents. No significant relationship was observed between sequence divergence and climatic niche overlap (y=2.063x + 0.247; *p*=0.5097, adj-R^2^=-0.161).

### Rates of multivariate climatic niche evolution in plant polyploid and diploid species

Our analyses found that polyploid species had faster rates of multivariate climatic niche evolution relative to congeneric diploids. Polyploid species consistently had significantly faster rates (*p*<0.05) under different model selection approaches (AICc and LRT), and across all four multivariate climatic niche traits of niche mean and breadth, except for the LRT model of PC2 for multivariate niche breadth (Table 2). In general, the AICc and LRT found similar qualitative results (Figure 3), however quantitative estimates of relative rates did differ (see Table S4). The following brief descriptions relate to results from the more conservative AICc models.

**Table 2.** Summary of multivariate niche mean and niche breadth analyses. Both models of corrected Akaike Information Criterion (AICc) and likelihood ratio test (LRT) displayed. Levels of significance displayed (*p*<0.01)** and (*p*<0.05)* are based on a two-tailed binomial exact test assuming an equal probability of each rate scenario (*p*=0.333).

| AICc Model Niche<br>Trait | Faster Rate in<br>Polyploids | No Rate<br>Difference | Lower Rate in<br>Polyploids |
| --- | --- | --- | --- |
| PC1 Mean | 22** | 4 | 7 |
| PC2 Mean | 26** | 2 | 5 |
| PC1 Breadth | 25** | 2 | 6 |
| PC2 Breadth | 19** | 3 | 11 |
| LRT Model Niche Trait | Faster Rate in<br>Polyploids | No Rate<br>Difference | Lower Rate in<br>Polyploids |
| PC1 Mean | 19** | 2 | 12 |
| PC2 Mean | 25** | 1 | 7 |
| PC1 Breadth | 26** | 1 | 6 |
| PC2 Breadth | 15 | 0 | 18 |

\*( $p < 0.05$ )
\*\*( $p < 0.01$ )

**Figure 3.**
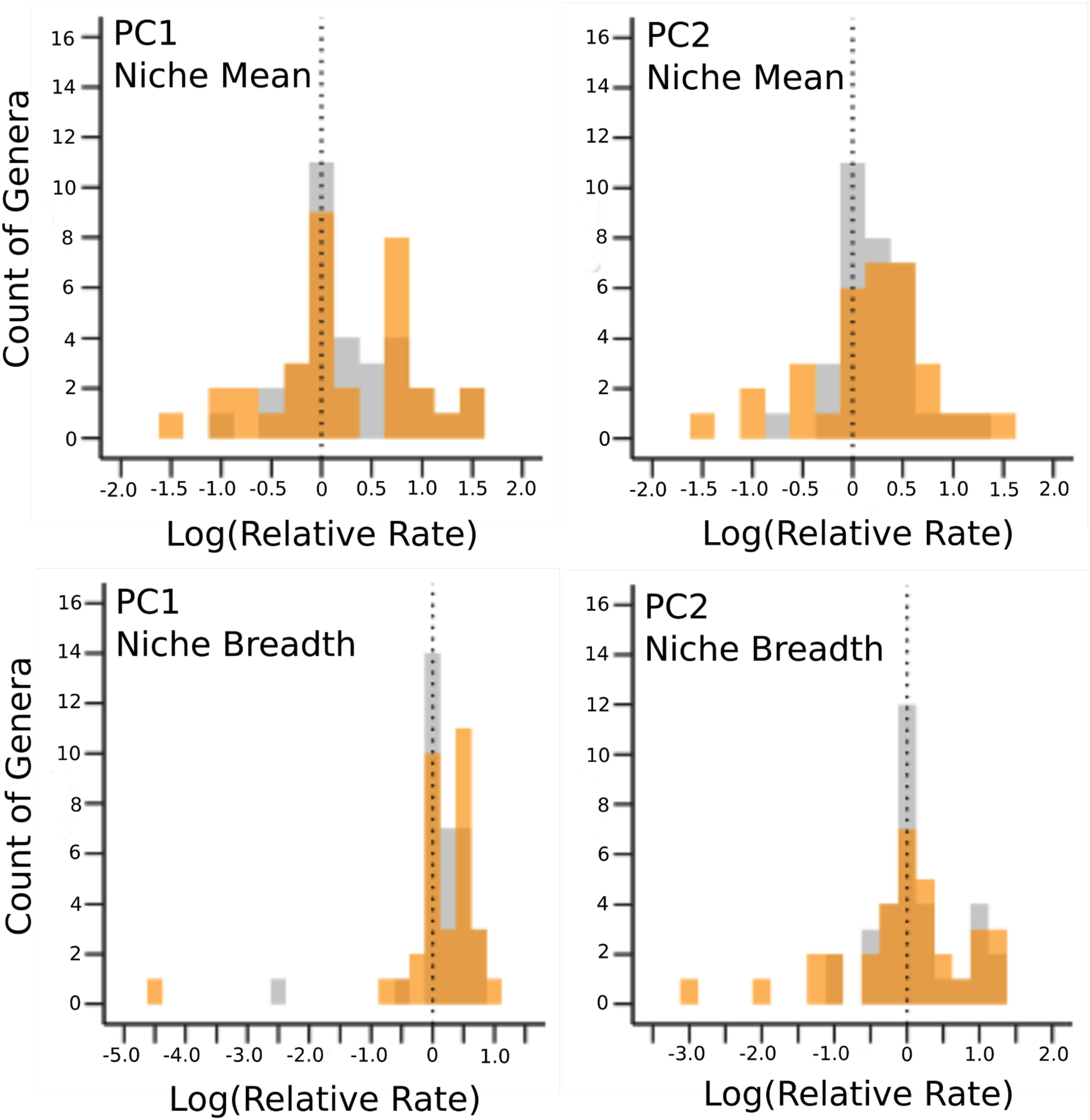
Relative rate estimates for multivariate climatic niche mean and breadth. The x-axis represents the log transformed relative rate difference between the polyploid species of a genus and their diploid congeners, and the y-axis represents the number of genera in each bin. The vertical dashed line signifies whether polyploids had a lower inferred rate (negative), faster inferred rate (positive), or no rate difference (neutral=0). Results from both the AICc (grey) and LRT (orange) models are displayed.

Across 22 of 33 genera, polyploid species had a higher estimated rate of change in the PC1 mean relative to diploid species. In contrast, polyploid species were slower than diploid species in the rate of change for PC1 mean in only seven genera. We found no significant difference in four genera. The lowest inferred rates of change of the PC1 mean for polyploid species was 0.10 times slower than diploids in *Centaurea*, whereas the highest rate was 38.89 times faster for polyploid species in *Aconitum*. Similarly, polyploid species had a higher rate of change relative to diploid species in the PC2 mean among 26 of 33 genera, whereas polyploid species were slower in five genera and not different than diploid species in two genera. For the PC2 mean, the inferred rates ranged from a low of 0.16 times slower for polyploid species in *Nicotiana* to a high of 14.39 times faster in *Dryopteris*.

Polyploid species also had more rapidly evolving multivariate niche breadths than their diploid relatives. Niche breadth represents the variance for each bioclim variable that a diploid or polyploid species experiences across its entire range in multivariate space. We found that the breadth of PC1 evolved faster in polyploid species in 25 of 33 genera, diploid species had higher rates in six genera, and there was no difference between polyploid and diploid species in only two genera. The inferred rates ranged from a low of 0.004 times slower for polyploid species in *Centaurea* to a high of 5.78 times faster in *Muhlenbergia*. Similarly, polyploid species in 19 of 33 genera had higher rates of evolution in niche breadth for PC2. Rates were lower for polyploid species in only eleven genera, and there was no difference in rate between polyploid and diploid species in three genera. The inferred rates of change of the PC1 breadth ranged from a low of 0.08 times slower for polyploid species in *Anemone* to a high of 22.92 faster in *Asplenium*.

Within a genus, similar rate trends were found for multivariate niche mean (PC1–2) and breadth (PC1–2) with few exceptions. For example in *Muhlenbergia*, polyploid species were estimated to have a 1.09 times higher rate than diploids for PC1 niche mean but a lower rate of 0.73 times the rate of diploids for PC2 niche mean. This variability in rates between PC1 and PC2 multivariate niche mean were found for nine genera (*Anemone, Centaurea, Eragrostis, Muhlenbergia, Orobanche, Poa, Primula, Silene,* and *Solanum*), and for PC1 and PC2 multivariate niche breadth 12 genera (*Aconitum, Anemone, Galium, Geranium, Nicotiana, Orobanche, Phacelia, Primula, Ranunculus, Silene, Solanum,* and *Veronica*). These include genera where there was no inferred difference between polyploid and diploid species on one PC axis but a difference was found on the other PC axis. Two genera had a more labile switch in whether polyploid species were inferred as having a higher or lower rate of multivariate niche evolution. The genus *Aconitum* had faster inferred multivariate niche mean rates for polyploid species in PC1 and PC2, but lower inferred rates of multivariate niche breadth in PC1 and PC2. Conversely, *Eragrostis* had lower inferred multivariate niche mean rates for polyploid species in PC1 and PC2, but higher inferred rates of niche breadth in PC1 and PC2. Except for the genera *Cuphea* and *Plantago* that had lower inferred rates for polyploid species across all niche traits, and *Fuchsia* which had no difference in rate between polyploid and diploid species for all four multivariate traits, all other genera had at least one multivariate niche trait with a greater inferred rate of evolution in polyploid species.

We used a simulation approach to assess if the correlated trends in the rates of multivariate niche traits could have been inferred by chance (see Table S5). Most genera had zero or only one axis with a rate direction that was indistinguishable from our null expectation (*p*>0.05). However, the genera *Draba, Ranunculus*, and *Veronica*—which had high proportions of polyploid species—also had the only instances where two or more axes were inferred to have a pattern similar to chance.

## Discussion

Polyploid speciation is one of the most common forms of sympatric speciation in plants. These species begin with substantial post-zygotic reproductive isolation, but there has been mixed support for the importance of ecological differentiation during polyploid speciation (Felber-Girard et al. 1996; Petit & Thompson 1999; Martin & Husband 2009; Glennon et al. 2014; Marchant et al. 2016). We found that the ecological niches of polyploid species often evolved faster than their diploid relatives across vascular plants. This was the case in the two different assessments of ecological niche differentiation—comparisons of niche overlap with diploid relatives and phylogenetic estimates of the rates of multivariate niche evolution. We found a majority of allopolyploid species from 54 trios had less climatic niche overlap with parents than the parents had with each other. Consistent with this observation, we also found that a majority of polyploid species from 33 genera had significantly faster rates of multivariate niche mean evolution as well as multivariate niche breadth evolution compared to their diploid relatives. These results indicate that ecological differentiation is a common and likely important component of polyploid speciation.

Our finding that polyploid species shift ecological niches faster than their diploid relatives is consistent with theoretical expectations. A significant ecological obstacle to polyploid establishment is minority cytotype disadvantage (Levin 1975). Nascent polyploid species are initially present at low numbers in populations of their diploid progenitors and must overcome frequency dependent gametic competition with their parents. Although regional coexistence may be possible without niche differentiation through stochastic processes and local dispersal (Baack 2005), simulations consistently find that polyploid establishment is promoted by ecological niche differentiation (Levin 1975; Fowler & Levin 1984; Felber 1991; Rodriguez 1996; Husband 2000; Rausch & Morgan 2005; Fowler & Levin 2016). Analyses of paleopolyploidy also indicate that polyploid species survived mass extinction events better than their diploid relatives (Vanneste et al. 2014; Lohaus & Van de Peer 2016). The relatively fast rates of ecological differentiation we observed may explain this phylogenetic pattern. Our results confirm that ecological divergence from progenitor species is a common and likely critical step in polyploid speciation. Polyploid species may also be differentiated from their progenitors in other dimensions, such as phenology or pollinators. Our current estimates should be considered a lower bound on the degree of ecological differentiation of polyploid and diploid species. Future work that leverages the growing body of trait data (Maitner et al. 2017) could extend our analyses of climatic niches to better capture the ecological divergence of polyploids.

The rapid niche shifts we observed in polyploid species may stem from the immediate phenotypic effects of WGD. Recently formed polyploid species manifest allometric phenotypic changes from an increase in nuclear DNA content due to positive relationships with cell size and volume (Speckman et al. 1965; Bennett 1972; Cavalier-Smith 1978; Melargno et al. 1993; Beaulieu et al. 2008; Chao et al. 2013). These effects on the phenotype are numerous, at times idiosyncratic, but important to how nascent polyploids interact with the abiotic and biotic environment. For example, Chao et al. (2013) found that the increase in cell size that accompanied WGD caused increased potassium uptake and salinity tolerance. This type of abrupt, genotype independent shift of ecological niche may explain some of the rapid shifts in ecological niche observed in our analyses. Other physiological differences associated with WGD include changes in propagule volume (Barrington et al. 1986), size and density of stomata (Sax & Sax 1937; Maherali et al. 2009), resistance to drought and cold (Levin 1983), and secondary metabolites and phenology (Levin 1983; Seagraves & Anneberg 2016). The many cases examined in our analyses provide a starting point for exploring the contribution of these potential avenues for WGD—independent of genotype—to alter plant physiology and ecological niche.

Polyploid species may also shift their ecological niches faster than their diploid relatives because of differences in genetic variation and selection associated with ploidal level increase. Evolutionary genetic theory predicts that polyploid species adapt faster than diploid species under certain circumstances (Orr & Otto 1994; Otto & Whitton 2000; Otto 2007). Recent experimental studies support these predictions in yeast (Selmecki et al. 2015). The capacity for a greater response to selection may stem from increased genetic diversity from WGD paralogs, although the proximate mechanisms are diverse. Immediately following WGD an increase in alleles may mask deleterious mutations and increase the probability of acquiring new beneficial mutations (Otto 2007). Paralogs can also diverge in function through sub– or neofunctionalization (Force et al. 1999; Lynch & Force 2000), which may lead to the evolution of novel adaptive traits (Levin 1983; Flagel & Wendel 2009; Edger et al. 2015). In addition, the increased genetic variation of polyploid species may come from rapid and diverse structural genomic changes (Song et al. 1995; Chester et al. 2012) and the multiple origins of such populations (Ownbey 1950; Werth et al. 1985; Brochmann et al. 1992; Doyle et al. 2003; Soltis & Soltis 1999). Recent analyses of how polyploids respond to abiotic stress (Bardil et al. 2011; Akama et al. 2014; Paape et al. 2016; Takahagi et al. 2018) suggest this variation is important for polyploid establishment and that it may play a role in the rapid climatic niche shifts we observed.

Previous analyses of niche differentiation in polyploid species found less consistent patterns (Felber-Girard et al. 1996; Petit & Thompson 1999; Martin & Husband 2009; Glennon et al. 2014; Marchant et al. 2016). Our consistent observation of faster rates of climatic niche evolution in polyploid species relative to diploids may be because of three main factors. One, we did not restrict our dataset of diploid and polyploid species to either a single clade or a single regional or continental area. This allowed us to expand our sample size to the largest dataset to date of polyploid and related diploid species. Second, our analyses took into account phylogeny and sequence divergence in relation to the change in climatic niche. Although previous analyses have investigated smaller datasets of auto-or allopolyploid species and their diploid progenitors (Glennon et al. 2014; Marchant et al. 2016), none examined a genus with the phylogeny taken into account. Finally, our dataset may be biased towards allopolyploids. Our first analysis focussed solely on these taxa. Auto-and allopolyploids are present, on average, in nearly equal proportions in nature (Barker et al. 2016c). However, autopolyploids are much less likely to be named than allopolyploids and probably under-represented in our analyses. Given that this bias is inherent in the taxonomy of polyploid species, most analyses of polyploid biogeography are likely impacted by it.

The hybrid origins of allopolyploid species may provide a significant source of increased genetic diversity compared to autopolyploid species. Hybridization can increase the amount of additive genetic variance which can be adaptive depending upon the environmental context (Anderson 1949; Stebbins 1959; Lewontin & Birch 1966; Stebbins 1985; Seehausen 2012; Bailey et al. 2013; Eroukhmanoff et al. 2013; Grant & Grant 2016). This genetic variance can also lead to a range of different phenotypes including intermediate, mosaic, or transgressive phenotypes (Rieseberg et al. 1999; Dittrich-Reed & Fitzpatrick 2013). Allopolyploid species inherit these novel evolutionary combinations, which may allow them to explore divergent ecological niches from their diploid progenitors. Indeed, recent evidence from natural populations of the *Alyssum montanum* species complex highlights the importance of hybridization in the evolution of climatic niches. Allopolyploid cytotypes of *A. montanum* have more divergent and higher rates of climatic niche evolution than their related autopolyploid cytotypes and diploid progenitors (Arrigo et al. 2016). Whether such biology is the norm for polyploid species remains to be tested, but our results indicate that it could be common across vascular plants.

A majority but not all allopolyploid species were faster in their rate of climatic niche evolution relative to their diploid progenitors or congeners. Such exceptions may be due to methodological limitations or a true lack of climatic niche evolution of these allopolyploid species. In the analyses of climatic niche overlap, we included all nineteen bioclimatic variables at 2.5 minute resolution which corresponds to roughly 21.62 km^2^ at the equator or 12.58 km^2^ at 40° latitude. Although this resolution increases with increasing latitude it may be too coarse to capture fine scale climatic niche differences (Kirchheimer et al. 2016). Additionally, the abiotic conditions of climate comprise a subset of the many axes that may be important in defining a taxon’s realized niche, and one set of abiotic axes that we did not consider are those related to edaphic conditions. Differences in edaphic tolerances have been shown to strongly differentiate closely related plant taxa (van der Niet & Johnson 2009; Anacker & Strauss 2014; Shimuzu-Inatsugi et al. 2017), but will not leave a signature of climatic niche evolution because they will appear to have the same climatic niche. This is especially true of edaphic endemics when the progenitor(s) are geographically more widespread, such as *Layia discoidea* (Gottlieb et al. 1985, Gottlieb 2004; Baldwin 2005), or populations on mine tailings as in *Anthoxanthum odoratum* (Antonovics et al. 1971, 2006). Differences in edaphic tolerances may explain some of the cases in our analysis where similar climatic niches were observed between allopolyploid species and their diploid progenitors.

The lack of an association of the pairwise sequence divergence of allopolyploid species and diploid progenitors and the amount of niche differentiation suggests that ecological differentiation may happen quickly upon formation. This is consistent with previous empirical observations of rapid phenotypic change (Ramsey & Schemske 2002). Alternatively, it may also reflect a lack of power with our available samples and molecular marker. We had relatively few data points for trios with pairwise sequence divergence and at this time we are cautious in our interpretations of such results but find it worthwhile for future study.

We also found that polyploid species consistently evolve niche breadths at faster rates than their diploid congeners, but we did not explicitly examine whether polyploid species had broader or narrower niche breadths. Understanding the role of niche breadth on nascent polyploid establishment is ripe for future study because niche breadth has a strong relationship to both speciation and extinction processes and ultimately diversification (Janzen 1967). Depending on the conceptual framework, both broad and narrow niche breadths may promote diversification (Sexton et al. 2017). Broad niche breadths may promote diversification because generalists are more likely to have larger ranges (Slayter et al. 2013) and thus lower extinction rates due to a possible ‘dead-end’ effect of specialization (Schluter 2000). However, species with narrow niche breadths have been found to have faster rates of niche evolution (Huey & Kingsolver 1993; Whitlock 1996; Fisher-Reid et al. 2012) as well as higher diversification rates (Hardy & Otto 2014; Rolland & Salamin 2016; Qiao et al. 2016). Future analyses that link changes in niche breadth to the genetic variation of polyploid species may contribute to our understanding of the macroevolutionary patterns of polyploid diversification.

Finally, our results provide insight into speciation processes more widely. A central question of speciation research is how intrinsic postzygotic reproductive isolation arises within populations of reproductively compatible individuals. Many models of speciation, such as the classic Bateson-Dobzhansky-Muller model (Bateson 1909; Dobzhansky 1934; Muller 1939), solve this problem by proposing that postzygotic reproductive isolation evolves after geographic or ecological isolation. Although speciation with gene flow is possible (Barluenga et al. 2006; Niemiller et al. 2008; Nosil et al. 2009; Yeaman & Otto 2011; Yeaman & Whitlock 2011; Feder et al. 2012; Nosil & Feder 2012; Martin et al. 2013; Wolf & Ellegren 2016; Samuk et al. 2017), most models require some degree of physical separation for intrinsic postzygotic isolation to arise (Coyne & Orr 2004; Gavrilets 2004). In contrast, polyploid species being with substantial postzygotic isolation from their progenitors while also sympatric with one or both parental species (Ramsey & Schemske 1998, 2002). By inverting the usual order of events during speciation, polyploid species provide a unique test of the importance of ecological differentiation to speciation in general. Our results affirm the importance of ecological differentiation to speciation, even in species that are founded with significant post-zygotic reproductive isolation.

## Acknowledgements

We would like to thank Z. Li, S.A. Jorgensen, X. Qi of the Barker Lab, and M.J. Sanderson, R. Ferrier, M. Worobey, R.H. Robichaux for comments on earlier drafts. Hosting infrastructure and services provided by the Biotechnology Computing Facility (BCF) at the University of Arizona. This research was supported by NSF-IOS-1339156 and NSF-EF-1550838.

